# Light transport modeling in highly complex tissues using implicit mesh-based Monte Carlo algorithm

**DOI:** 10.1101/2020.10.11.335232

**Authors:** Yaoshen Yuan, Shijie Yan, Qianqian Fang

**Author notes:** http://fanglab.org/.

## Abstract

The mesh-based Monte Carlo (MMC) technique has grown tremendously since its initial publication nearly a decade ago. It is now recognized as one of the most accurate Monte Carlo (MC) methods, providing accurate reference solutions for the development of novel biophotonics techniques. In this work, we aim to further advance MMC to address a major challenge in biophotonics modeling, i.e. light transport within highly complex tissues, such as dense microvascular networks, porous media and multi-scale tissue structures. Although the current MMC framework is capable of simulating light propagation in such media given its generality, the run-time and memory usage grow rapidly with increasing media complexity and size. This greatly limits our capability to explore complex and multi-scale tissue structures. Here, we propose a highly efficient implicit mesh-based Monte Carlo (iMMC) method that incorporates both mesh- and shape-based tissue representations to create highly complex yet memory efficient light transport simulations. We demonstrate that iMMC is capable of providing accurate solutions for dense vessel networks and porous tissues while reducing memory usage by greater than a hundred- or even thousand-fold. In a sample network of microvasculature, the reduced shape complexity results in nearly 3x speed acceleration. The proposed algorithm is now available in our open-source MMC software at http://mcx.space/#mmc.

## 1. Introduction

The Monte Carlo (MC) method is widely used as the gold standard for modeling light propagation in complex media such as human tissues [1,2]. Due to great efforts in accelerating MC methods using emerging parallel hardware, such as graphics processing units (GPUs), in the past decade, MC methods have thrived in the field of biomedical optics [3–6], benefiting a broad range of applications such as functional brain imaging [7], photobiomodulation treatment planning [8,9], new instrument prototyping [10], food and agriculture [11], and marine science [12], among others. Earlier MC simulation methods [1] can handle relatively simple geometries, such as layered media. The added capability to model 3-D voxel-based heterogeneous domains [3,13–15] represents a major milestone towards broader utilities. However, using voxels to represent shapes with curved boundaries leads to expensive memory and computational costs because high voxel densities are usually needed to retain boundary accuracy.

About 10 years ago, we published one of the first mesh-based MC (MMC) algorithms in the inaugural issue of this journal [16]. The MMC method is designed to overcome the limitations of voxel-based MC by performing ray-tracing computations in tetrahedral meshes. Because tetrahedral meshes present greater flexibility in representing arbitrary shapes, MMC can model light propagation in complex anatomical structures with high accuracy and memory efficiency. Noticeable improvements in simulation accuracy over voxel-based MC have been reported in various studies [16–18].

Since the initial proposal of the MMC algorithm, there has been tremendous growth in developing refined and accelerated MMC methods to address various challenges in the field. A number of works were targeted towards accelerating MMC by utilizing parallel architectures, including the extensions to single-instruction multiple-data (SIMD) [19] and advanced vector extensions (AVX) SIMD instructions [20] on the CPU, as well as the use of NVIDIA CUDA [5] and OpenCL [6, 21] to gain substantial acceleration through modern GPUs. New efforts have also led to the generalization of MMC to support wide-field illumination and camera-based detection [22,23], as well as a dual-grid MMC (DMMC) approach combining a coarsely tessellated mesh for ray-tracing with a fine voxelated domain for storage [17]. Other new MMC-based approaches have been proposed to address challenges in ease-of-use [24], modeling anisotropic tissues [21], studying new imaging techniques [25], and solving inverse problems [26,27].

Despite exciting advances thus far, the standard for MC light modeling is constantly being challenged as biophotonic techniques continue their endeavors towards probing deeper and increasingly complex and multi-scaled tissues. In order to facilitate the rapid growth in emerging optical imaging techniques and offer a better understanding of tissue physiology at all lengthscales, increasingly higher demands have been placed upon efficient computational methods and light-tissue interaction modeling. For example, in fluorescence laminar optical tomography (FLOT) – a mesoscopic imaging technique, MC method was found for characterizing light-tissue interaction in brain cortex [28]. In optical coherence tomography (OCT) – a modality with a resolution of a few micrometers – MC methods have been used for computing the angiography images [29]. On the microscopic scales, multi-photon microscopy continues pushing limits in high spatial resolution and researchers have been using MC for improving the penetration depth [30,31]. Aside from numerous studies using MC for processing functional near-infrared spectroscopy (fNIRS) data [18] – a macroscopic imaging technique for neural imaging, a recent study has attempted to build connections between fNIRS hemodynamic signals using MC simulations with a simplified mesh-based vessel network [32]. From these recent studies, it is strongly evident that an accurate, fast and versatile MC model could offer widespread impact towards pushing frontiers in biophotonics research.

We want to point out that, compared to layfer- and voxel-based MCs, MMC is already a significant step forward in versatility and accuracy, and is capable of modeling arbitrarily complex domains. However, creating accurate mesh models from such domains often leads to extremely dense meshes with large node and element numbers. This could have a profound impact on simulation speed, maximum domain volumes, as well as output resolution. Using increasingly sophisticated mesh models can quickly become a bottleneck on memory-limited devices such as GPUs [6]. While implicit shape-based MC techniques have previously been explored [33–36], their utilities have largely been limited to domains with simple geometries, lacking the complexity necessary to meet the needs of multi-scaled tissue simulations, such as dense vessel networks derived from microscopic imaging [37]. Therefore, further extensions that enable MMC to produce more efficient and robust simulations in such scenarios are highly desirable.

Here, we propose a highly memory-efficient and versatile MMC algorithm – implicit MMC or iMMC – that combines shape-based modeling with mesh-based MC, permitting efficient simulations in large-scale complex tissue structures. In this approach, only a coarse “skeletal” tetrahedral mesh is needed to represent essential shapes, such as the center-lines of blood vessels or centroids of pores or cavities, while detailed tissue structures are represented via implicitly defined shapes associated with the selected edges, faces and vertices of the skeletal mesh. Because of this hybrid shape representation, iMMC is capable of modeling extremely complex and multi-scaled tissues, such as dense networks of microvasculature or microporous alveolar lung tissue, using only fractional nodes/elements when compared to conventional MMC.

In the remainder of this paper, the hybrid shape-mesh modeling and the corresponding mesh generation procedures are introduced in Section 2. In Section 3, we validate the iMMC algorithm by comparing it with conventional MMC [16] and dual-grid MMC [17] methods in several benchmarks. A real-world example, in which iMMC is applied to model a highly complex microvascular network, is also reported in Section 3. Finally, in Section 4, we quantify the improvement in memory utility and simulation speed, and discuss future directions.

## 2. Methods

Unlike the traditional MMC approach which relies on tetrahedral meshes to explicitly represent structures, the iMMC algorithm implicitly defines complex tissue domains by attaching parameter-based shapes onto the edges, nodes and faces of a tetrahedral mesh. In the following subsections, we will separately describe our processing pipelines for 1) edge-based iMMC (e-iMMC), where implicitly defined cylindrical domains are associated with selected edges to represent vessel/airway networks, 2) node-based iMMC (n-iMMC), where implicitly defined cavities are associated with vertices for modeling porous media, and 3) face-based iMMC (f-iMMC), where a thin membrane is associated with the selected triangular faces of tetrahedral elements to model ultra-thin tissue layers such as human epidermis.

### 2.1. Edge-based iMMC

Let us first consider a complex vessel network [37] (also applicable to other tree-like tubular tissue structures, including airways in the lung, milk-ducts in the breast or collecting ducts in the kidney). To capture the overall shapes of such structures, we can separate the geometric information into two parts: the branch-like center-lines of the vessel network and the associated radius for each segment or branch of the center-line. To avoid creating high node densities when meshing thin-tube structures, as is the case in standard MMC, we only need to create a coarse “skeletal” mesh in iMMC, made from the Delauney tessellation of center-line nodes and those along the exterior boundaries while suppressing the addition of Steiner nodes [38]. Once the skeletal mesh is created, the center-line segments, now edges in a subset of tetrahedra, are subsequently labeled and attached with the corresponding radii of vessels, as obtained from image processing/segmentation tools. We want to note here that vessel segmentation is not the focus of this work and one should consider using appropriate pre-processing tools for obtaining vessel information before performing iMMC simulations.

To handle branching, more than one local edge of a tetrahedron can be labeled as vessels in an e-iMMC simulation. To simplify the computation in our current implementation, we support up to two edges per element for vessel modeling; higher order junctions involving more than 2 branches can be handled by refining the element next to the bifurcation until each element contains two or fewer branches. In Fig. 1(a), we illustrate a vessel-bearing element, where edge 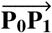 is associated with a vessel of radius *r*.

**Fig. 1.**
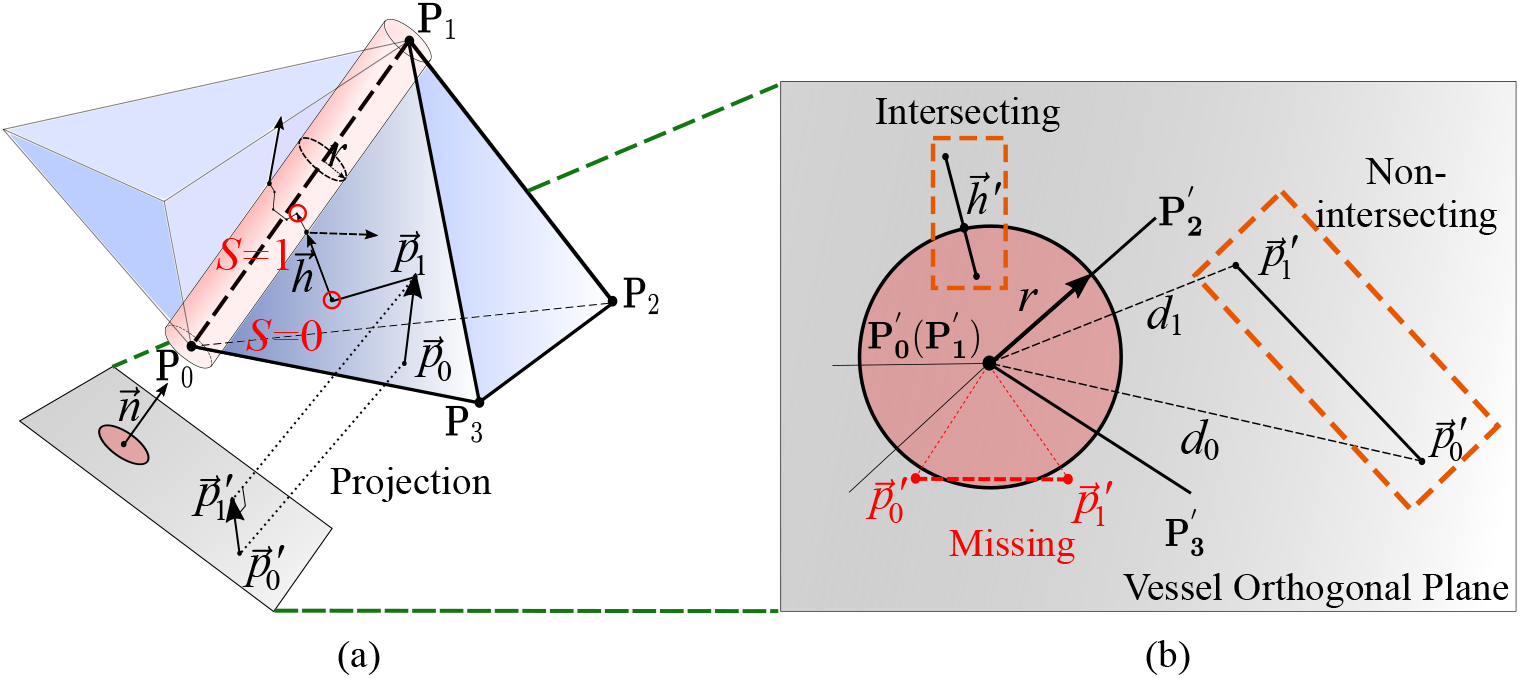
Illustrations of edge-based implicit mesh-based Monte Carlo (e-iMMC) algorithm. Here we show (a) the photon packet ray-tracing in a tetrahedral element with a labeled edge as the center line of a vessel segment (red), and (b) the ray-cylinder intersection testing along the orthogonal plane of the vessel. The dashed line boxes in (b) show examples of intersecting and non-intersecting rays.

The ray-tracing process in an e-iMMC simulation is largely similar to those in standard MMC or DMMC – a photon packet traverses between tetrahedra by performing ray-tetrahedron intersection testing and accumulates energy losses along its path [6,16]. The only difference is that when a packet enters a vessel-bearing element, intersections between its trajectories and implicitly-defined tubular vessel surfaces must be computed for each propagation step. As a result, an additional state (*S*) is recorded for each simulated packet to denote whether it is located inside a vessel (*S* = 1) or outside (*S* = 0), and is updated for each step. Using the *S* flag, one can then select the corresponding optical properties for scattering or absorption calculations.

Determining whether a photon’s path segment intersects with the cylindrical vessel surface is an essential part of the e-iMMC computation. For a given packet path segment within a vessel-bearing element, denoted as 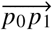 in Fig. 1, where edge 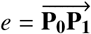 is designated as a vessel center-line with radius *r*, we first compute the distance between the starting point 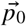 and vessel center-line *e* as *d*_0_ and between the end point 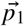 and *e* as *d*_1_.

As illustrated in Fig. 1(b), in the e-iMMC algorithm, we can perform a rigorous ray-cylinder intersection computation along each propagation step of the packet and determine 1) whether the path segment intersects with the cylinder, and 2) the entry and exit positions along the cylinder if an intersection occurs. Specifically, this can be done by solving the quadratic line-circle equation along the 2-D plane orthogonal to the vessel edge *e*. When an intersection is detected, the intersection points can be computed in the 2-D plane, denoted as 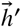 in Fig. 1(b), first and then mapped to the 3-D space, denoted as 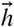 in Fig. 1(a).

We want to note that the above per-step testing provides good accuracy, but requires solving a quadratic equation at every step. To accelerate the computation, our current implementation utilizes a necessary but insufficient condition for the path segment to intersect with the cylinder:

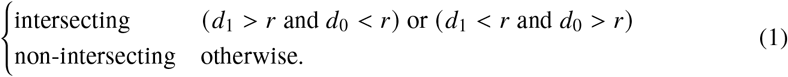

As shown in the projection view, Fig. 1(b), along the orthogonal plane to the vessel *e*, the above condition correctly determines the top two cases (dash-line boxes), but misses the bottom case (a false negative), where both 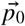 and 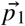 are located outside of the vessel. As demonstrated in our benchmarks in the Results section, the errors due to these false-negative tests are negligible for the tested domain shapes and optical properties. We want to point out that the false-negative cases may become more frequent when the average path lengths (related to both the mean-free-path and the average edge-length of the tetrahedron) of the packet is comparable or much larger than the vessel’s diameter. In such a case, one may choose to perform the rigorous ray-cylinder intersection testing or, alternatively, inflate the vessel radius to approximately compensate for the missing intersections. The inflation factor should be determined analytically or numerically.

When an intersection is detected, the packet is moved from 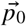 to the intersection point 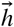. Whether a packet reflects from or transmits through the surface is then determined according to Fresnel’s law using the refractive indices of the vessel and the background tissue, similar to the reflection computation that occurs at triangular faces denoting tissue boundaries. The flag *S* is toggled after every transmission across the cylindrical boundary. The handling of energy deposition in iMMC is identical to that of the DMMC method [17], i.e. saving along a fine voxelated grid.

To handle the junction between two adjacent vessel segments, we add a sphere at each node along the vessel center-line, and apply the node-based iMMC method, detailed in the below section, to improve vessel continuity. The radius of the sphere is determined by max(*r*_1_, *r*_2_) where *r*_1_ and *r*_2_ are the radii of the two vessel segments.

### 2.2. Node-based iMMC and face-based iMMC

Sophisticated photon migration simulations may also arise when studying porous media, such as lung tissues, trabecular bones, fat emulsions, solutions with micro-bubbles, etc, as well as multi-scaled tissues, such as human epidermis, brain meninges, or cell membranes. Extending this hybrid shape and mesh based anatomical modeling to handle these types of complex media may foster a greater understanding of these complex systems by enabling more realistic and large-volume simulations. For these purposes, we have developed a node-based iMMC (n-iMMC) algorithm suitable for simulating porous domains by associating a spherical inclusion at mesh vertices and a face-based iMMC (f-iMMC) algorithm suitable for simulating thin membranes in a complex domain.

To demonstrate the basic utility of n-iMMC, in Fig. 2(a), we show one of the simplest porous domain simulations by defining spherical inclusions at a selected subset of mesh nodes. When simulating a porous domain with spherical inclusions (or cavities), the centroid and radius of each previously segmented inclusion (or generated synthetic data) are first extracted and recorded. Instead of creating spherical surface meshes for each inclusion, in n-iMMC, we create a very coarse tetrahedral mesh by simply tessellating the centroid positions along with other non-inclusion boundary nodes within the background media. We label each centroid node with the corresponding radius and tissue type. The radii may be uniform or follow a distribution. Any tetrahedral element that shares a centroid node is labeled as an “inclusion-bearing” element.

**Fig. 2.**
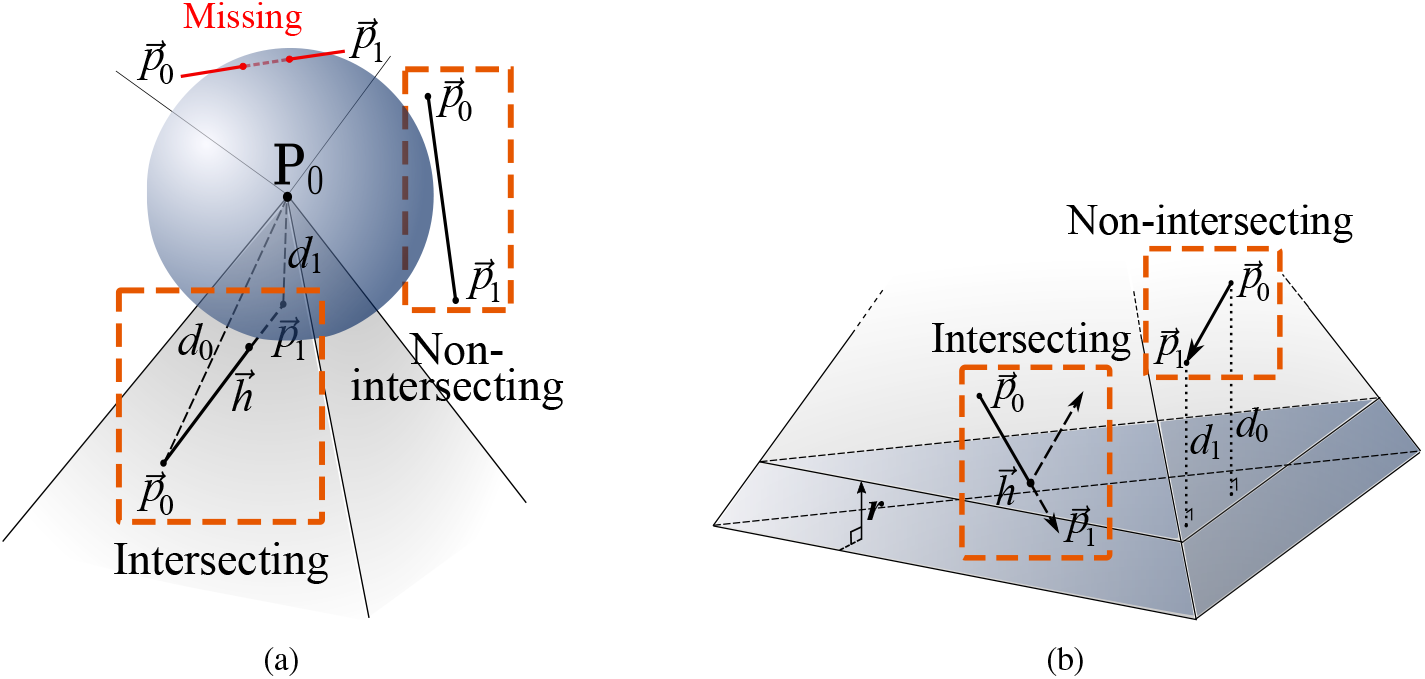
Illustrations explaining (a) the node-based implicit mesh-based Monte Carlo (n-iMMC) method and (b) face-based implicit mesh-based Monte Carlo (f-iMMC) method. The orange boxes highlight photon rays that either intersect or pass by the feature object.

Similar to the e-iMMC case discussed above, a photon packet that traverses a non-inclusion-bearing element follows steps identical to a standard MMC or DMMC simulation. However, when a packet traverses through an inclusion-bearing element, we have to perform an extra computation to determine whether the path segment intersects with the inclusion. For a path segment 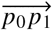 intersecting with a tetrahedral element that contains a spherical inclusion centered at *P*_0_ with radius *r*, the distances from the start point *p*_0_ and end-point *p*_1_ to the centroid node *P*_0_ are computed, denoted as *d*_0_ and *d*_1_, respectively.

Also similar to the aforementioned e-iMMC, one may choose to either perform a more rigorous ray-sphere intersection testing after each propagation step or apply Condition 1 to approximate and simplify the computation. When an intersection is detected, the intersection position 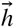 is then computed by solving the quadratic equations for line-sphere intersection. The photon is then moved to the point of intersection to evaluate for reflection versus transmission. When more than one inclusions are associated with an element, the intersection calculations should loop over each inclusion-containing vertex and the packet should be moved to the closest intersection position.

The face-based iMMC, or f-iMMC, allows to model thin-wall materials by defining a thickness, also denoted as *r* for simplicity, for each selected triangular face designated as part of a membrane. Modeling such a thin-membrane structure in a traditional MMC simulation often requires a very high number of elements because of the drastic difference in length scales between the membrane and background tissue. However, for f-iMMC, we only need to discretize the membrane outer surface, without considering its thickness, and will subsequently incorporate the thickness of the structure as an implicit parameter. This allows us to model arbitrarily thin membranes accurately without the worry of mesh density restrictions.

When a photon packet traverses inside a membrane-bearing element, the distances between the packet and the membrane faces will be computed during each propagation step to test whether it enters or exits the thin layer. Because the layer is flat, Condition 1 described above becomes both necessary and sufficient and does not result in any false tests. However, special scenarios must be considered when a photon is near the edges or corners of the element. See the following section for details.

### 2.3. Mesh generation and labeling

From the aforementioned subsections, one can see that one of the major differences between iMMC and conventional MMC is in mesh generation. An iMMC mesh is only a “skeletal” mesh that contains a coarse Delaunay tessellation of the essential geometric data – vessel center-lines, inclusion centroids, or face patches; the detailed structure shapes are now converted as implicit parameters and precisely tested in the photon ray-tracing calculations without needing to discretize. This not only results in dramatic reductions in mesh complexity and node/element counts, but also enhances accuracy by reinforcing shape boundaries as implicit parameters.

The creation of the coarsely tessellated tetrahedral mesh for iMMC is similar to the steps used for creating the coarse DMMC meshes [17], with the exception that the input data is not the triangular surfaces of the tissues, but the vessel center-lines (for e-iMMC), the pore centroids (for n-iMMC) or the membrane surface (for f-iMMC); these can be combined with triangular surface meshes representing non-multi-scale features, including bounding box or other tissue boundaries, similar to those used in standard MMC or DMMC. The feature edges, nodes, and triangular patches are recorded in a polyhedral file as the input for the open-source “TetGen” mesh generator [38]. To suppress the addition of Steiner points for mesh refinement, we used the “−Y” flag with TetGen. This results in tetrahedral meshes of the least element number without losing input geometric accuracy.

Subsequent data analysis is performed in MATLAB (MathWorks, Natick, MA, USA). Once the mesh has been generated, we find each of the feature line segments, nodes, or faces and create a new set of mesh input files in which the feature edges/nodes/faces are labeled with parameters to represent their cylindrical radii, spherical radii or layer thicknesses, respectively.

For illustration purposes, in Fig. 3, we show 3 simple mesh examples for edge-, node- and face-based iMMC, respectively. The vessel-mimicking edge in Fig. 3(a), inclusion-mimicking node in Fig. 3(b) and membrane-mimicking face in Fig. 3(c) are marked in red, and the feature-bearing tetrahedral elements are marked in light blue.

**Fig. 3.**
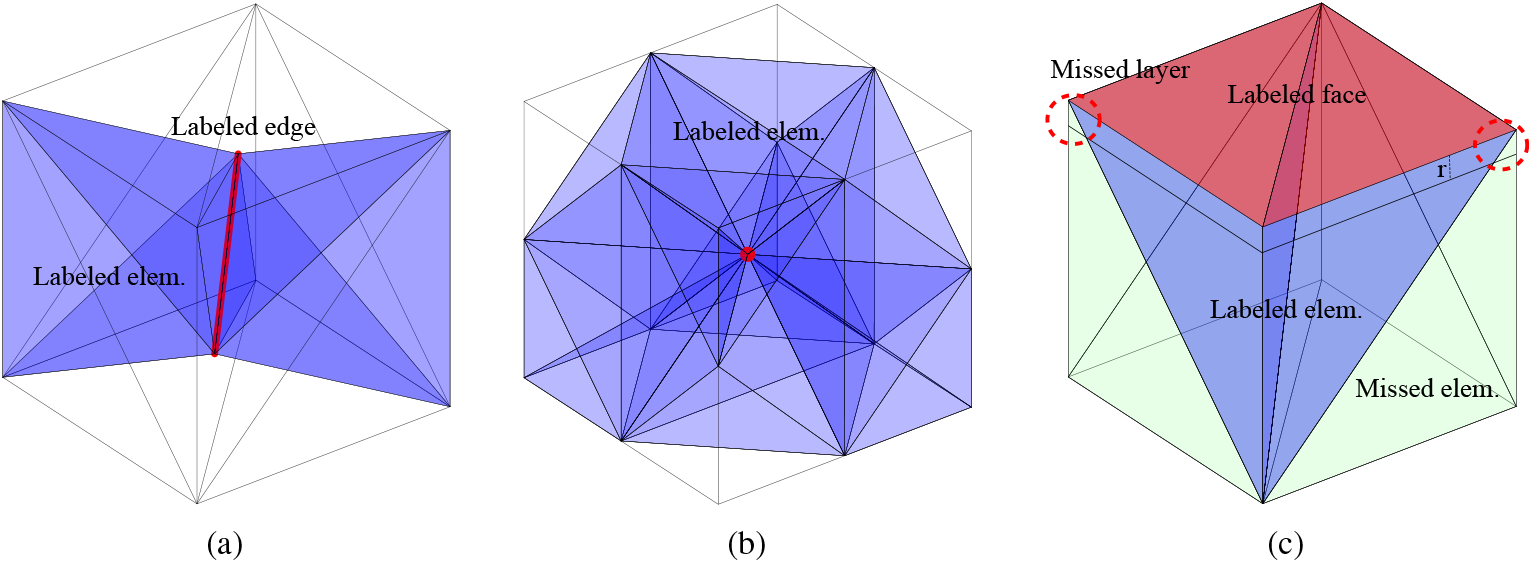
Illustrations of mesh generation and labeling for (a) edge-based iMMC, (b) node-based iMMC and (c) face-based iMMC. The labeled edge, node and face are indicated by red color and the feature-bearing elements are marked in blue. In (c), the red dashed circles indicate feature regions that are located inside the immediately neighboring elements (light green). The feature-bearing elements (blue) are “referred” by the neighboring elements when a photon traverses the latter.

We want to particularly mention that to properly account for all regions within the features (vessels or membranes) in e-iMMC and f-iMMC, we need to label not only those elements that directly contain edge/face features, but also those in their immediate vicinity. For example, in Fig. 3(c), a small part of the virtual layer (red dashed circles) are not included in the feature-bearing elements (light blue), but rather in the first-level neighboring elements that share nodes with the feature surface (light green). Labeling both types of elements allows f-iMMC to correctly compute light distributions among all regions. Similarly, as shown in light-green in Fig. 4(c), a small portion of the implicit vessel domain near one of the ends is inside neighboring elements and outside of the added spherical junctions. Therefore, labeling these adjacent elements and perform e-iMMC tests may lead to improved accuracy.

**Fig. 4.**
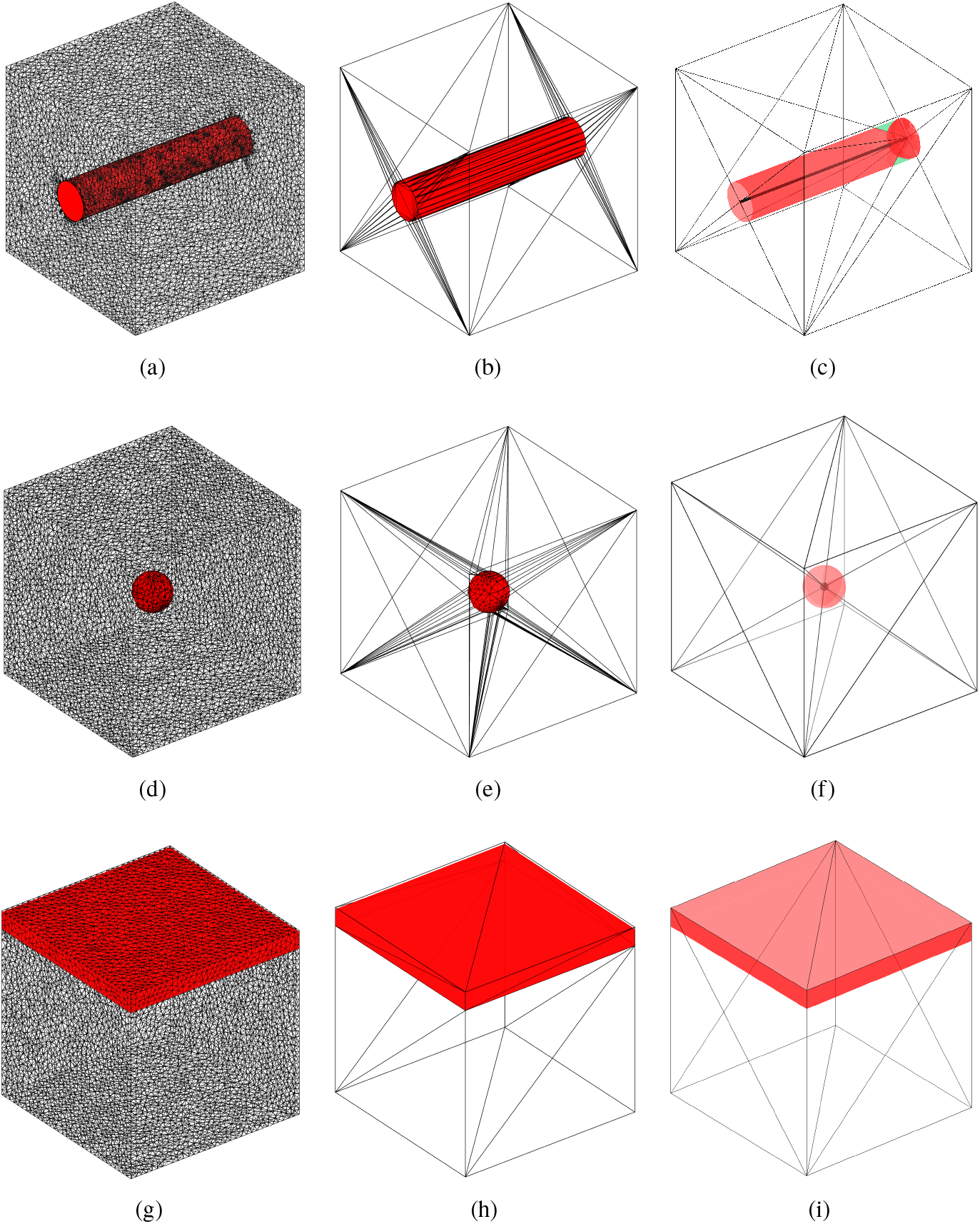
The meshes generated for three benchmarks and three Monte Carlo methods. Benchmarks (a-c) **B1**, (d-f) **B2** and (g-i) **B3** are built to validate the edge-based, node-based and and face-based iMMC algorithms, respectively. Each column, from left to right, shows the meshes created for standard mesh-based Monte Carlo (MMC), dual-grid mesh-based Monte Carlo (DMMC) and iMMC, respectively. The green shade in (c) indicates vessel regions within the elements adjacent to a vessel-bearing element.

## 3. Results

In this section, we first show a few simple benchmarks to demonstrate the essential features for the edge-, node- and face-based iMMC. Using these simple domains, we also validate iMMC by comparing it with conventional MMC and DMMC. Then, we show one real-world example by applying iMMC towards modeling photon transport within a highly complex microvascular network containing thousands of vessel branches.

### 3.1. Validation of iMMC methods

In Fig. 4, we show three benchmarks (referred to as **B1, B2** and **B3**) to validate e-iMMC, n-iMMC and f-iMMC, respectively. The domain sizes for all three benchmarks are 1 × 1 × 1mm^3^. A cylindrical vessel [Figs. 4(a)–4(c)], a spherical inclusion [Figs. 4(d)–4(f)] and a skin-like layered medium [Figs. 4(g)–4(i)] with a radius/thickness of 0.1 mm are placed in the domain. Three sets of meshes generated using either MMC, DMMC or iMMC are shown in the left, middle and right columns, respectively. Note that in Fig. 4(c), one sphere with the same radius of 0.1 mm is added at one end of the cylinder to mitigate the under-representation in e-iMMC caused by the tetrahedron geometry. The background medium has absorption coefficient *μ_a_* = 0.0458 /mm, scattering coefficient *μ_s_* = 35.65 /mm, anisotropy *g* = 0.9, and refractive index *n* = 1.37; those for the inclusions are *μ_a_* = 23.0543 /mm, *μ_s_* = 9.3985 /mm, *g* = 0.9, and *n* = 1.37, mimicking absorbers [14]. A downward-pointing pencil beam source is positioned at (0.5, 0.5, 1) mm. A total of 10^9^ photons are simulated in all cases. All simulations are executed on an AMD Ryzen Threadripper 3990x 64-Core CPU processor.

To quantify the memory-efficiency of iMMC, we list the numbers of nodes and elements for MMC, DMMC and iMMC for each benchmark in Table. 1. The simulation speeds (photons/ms) for each domain are also provided in the same table. All speeds and run-times in the paper are averaged from three repeated simulations.

**Table 1.**
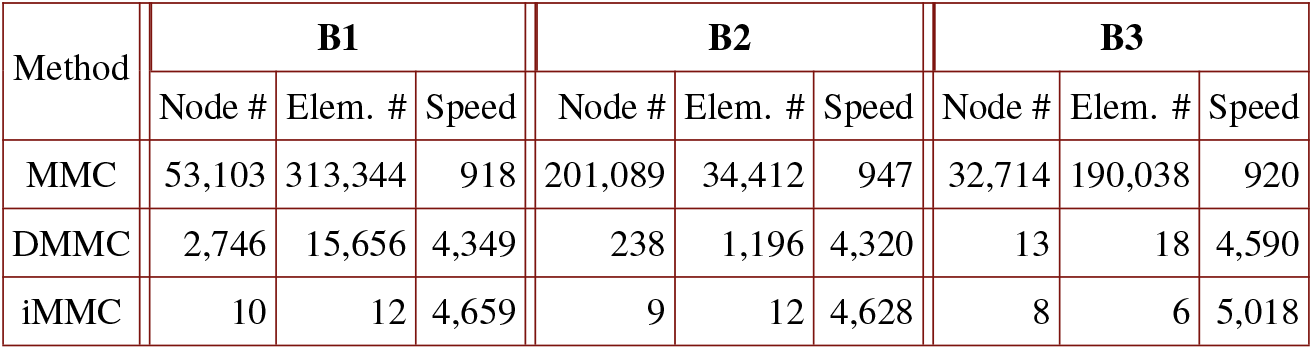
Comparisons between mesh-based Monte Carlo (MMC), dual-grid mesh-based Monte Carlo (DMMC) and implicit mesh-based Monte Carlo (iMMC) for three benchmarks – **B1, B2** and **B3**. The numbers of nodes (node #) and elements (elem. #) are compared below. The speeds (photons/ms) are also summarized.

To validate the accuracy of iMMC, in Fig. 5, we show cross-sectional plots of log-scaled fluence (J · mm^−2^) along the source planes to compare solutions derived from MMC, DMMC, and iMMC for the B1, B2, and B3 benchmark domains, respectively.

**Fig. 5.**
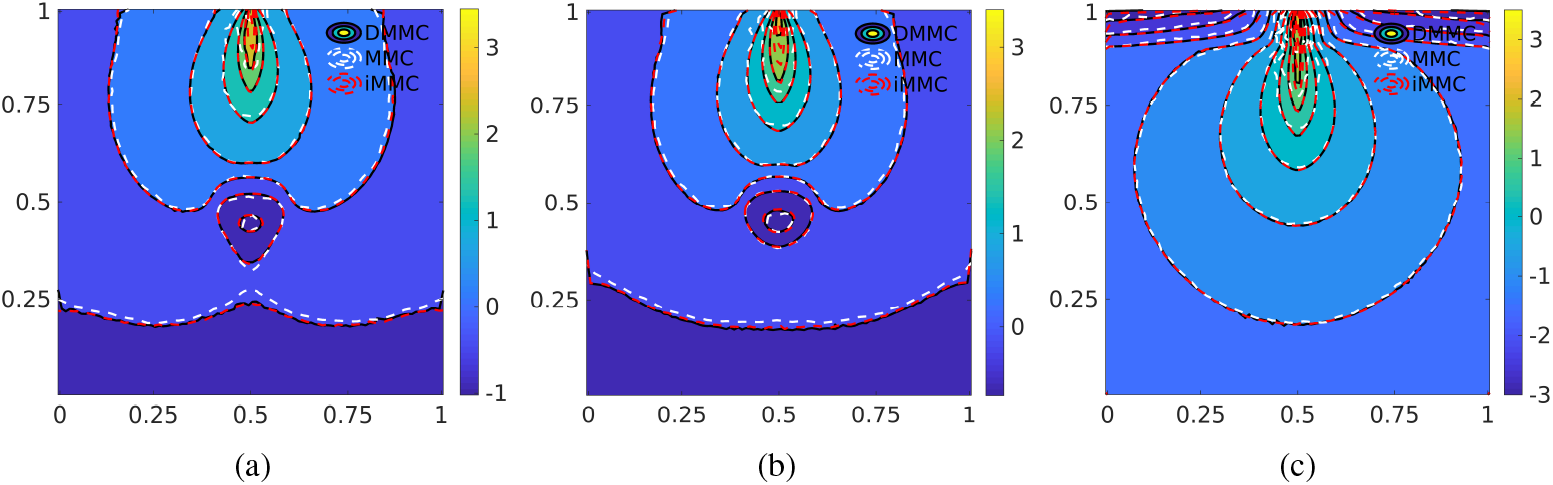
Log-scaled fluence (J · mm ^−2^) cross-sectional plots comparing three mesh-based Monte Carlo methods in three benchmarks (a) **B1**, (b) **B2** and (c) **B3**.

### 3.2. Application of iMMC in complex vessel networks

To show that iMMC simulations can be easily scaled to model life-sized complex tissue structures, we compare simulations performed using iMMC, voxel-based MC [3] and DMMC [17] within a 3-D synthetic microvascular network, shown in Fig. 6(a), created from data published in [39]. In Figs. 6(b)–6(d), we show zoomed-in views of three different domain representations created for 1) iMMC: a skeletal coarse mesh made of vessel center-lines with implicit parameters representing cylindrical vessels (red shade), 2) voxel-based MC: a rasterized microvascular network volume derived from the center-lines in 1), resulting in a volume of 326 × 305 × 611 voxels with isotropic 1 × 1 × 1 *μ*m^3^ voxels, and 3) DMMC: a tetrahedral mesh generated by first extracting a triangular surface from the above rasterized domain using the Iso2Mesh toolbox [40] and then creating a coarse tessellation [17] using TetGen [38]. In all three domain settings, a pencil beam light source is placed on the top at (150.5, 150.5, 611) μm pointing downwards. The blue circles in Figs. 6(b)–6(d) highlight some of the subtle differences between three discretization methods.

**Fig. 6.**
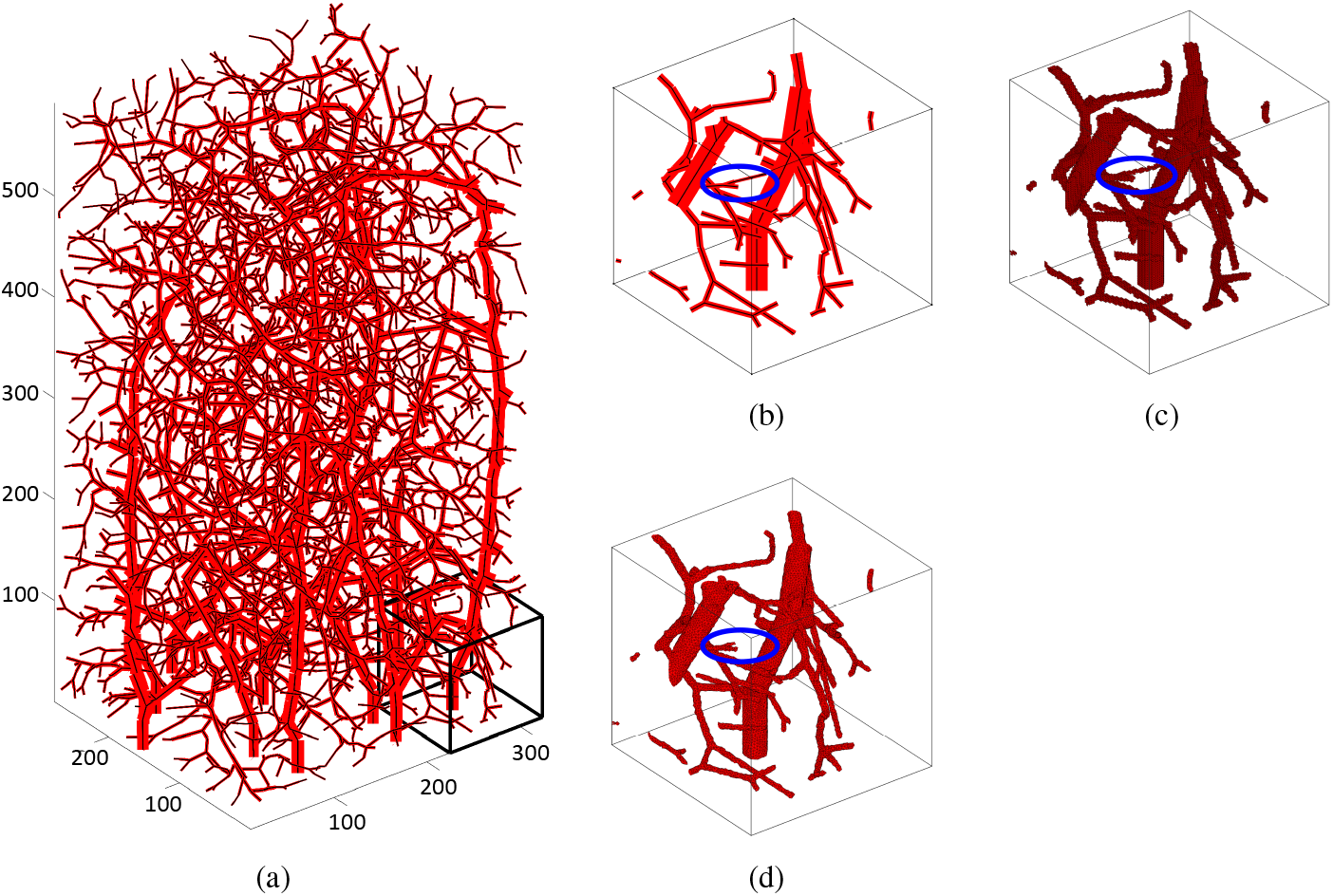
(a) The example domain (326 ×305 ×611 *μ*m^3^) of microvascular networks used in Monte Carlo modeling. A 100×100×100 *μ*m^3^ cubic region at the bottom right corner of the domain is magnified on the right for (b) implicit mesh-based Monte Carlo (iMMC), (c) voxel-based MC and (d) dual-grid mesh-based Monte Carlo. The black line in (a) and (b) represents the center-line of the vessels. The blue circles in (b)-(d) highlight a 1-voxel-wide vessel that is missing from the DMMC mesh due to limited meshing resolution.

In Figs. 7(a-c), to highlight the vessel geometry, we show the 3D cross-sectional plots of energy deposition per voxel (J) ranging from 0 to 2.5 × 10^−7^ J without log-scale at *y* = 150 *μ*m for iMMC, voxel-based MC using Monte Carlo eXtreme (MCX [3], http://mcx.space) and MMC, respectively. The red rectangle in Figs. 7(a-c) are further compared in Fig. 7(d) as contour plots (in log-10 scales).

**Fig. 7.**
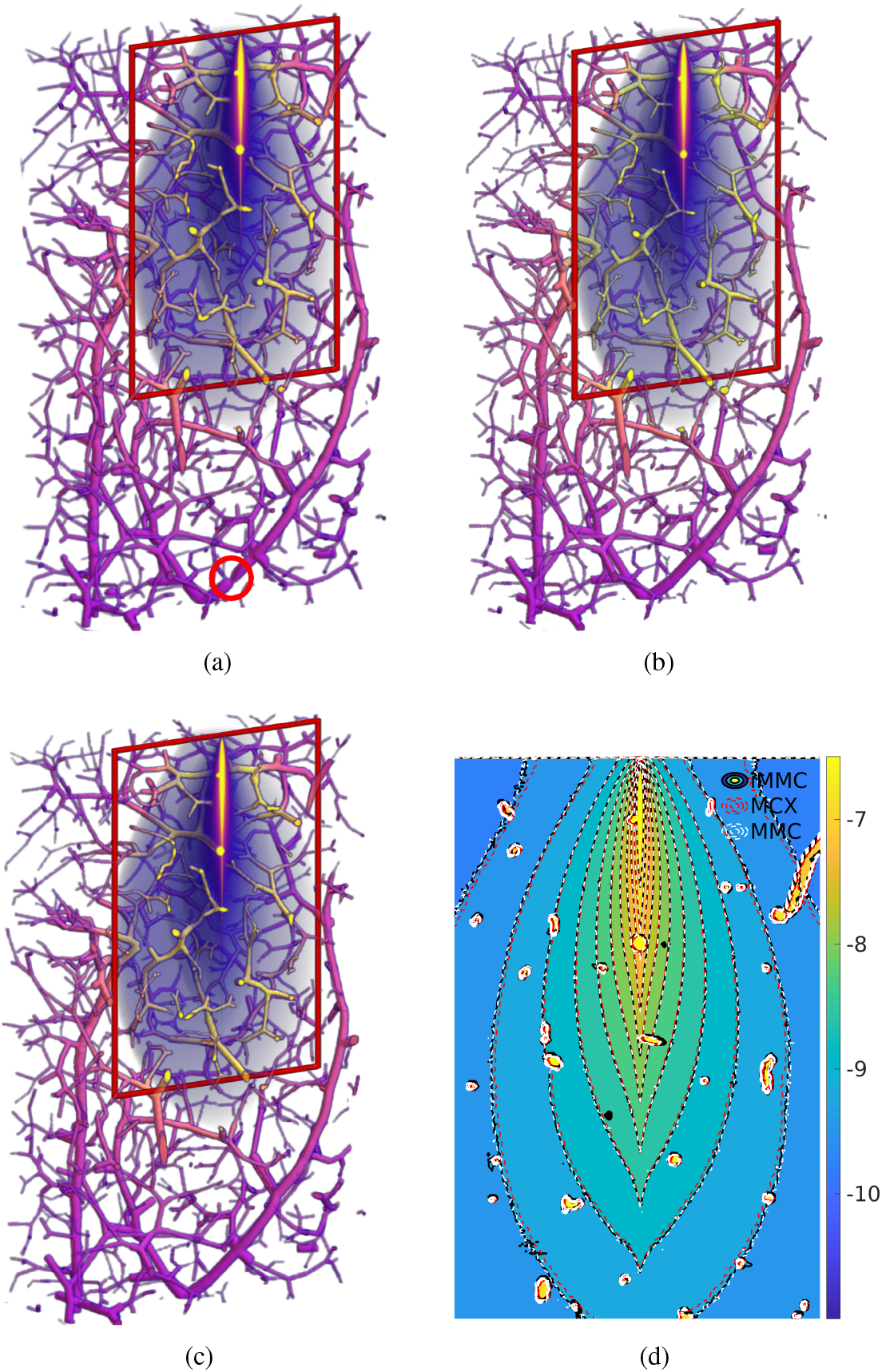
Three-dimensional plots of energy deposition per voxel (J) within a highly complex microvascular network using (a) implicit mesh-based Monte Carlo (iMMC), also shown in Visualization 1, (b) voxel-based Monte Carlo (simulated using MCX) and (c) dual-grid mesh-based Monte Carlo method (DMMC). The red circle in (a) indicates an artifact caused by approximations without considering adjacent elements next to a vessel element (see Fig. 4(c)). The contour plots of the fluence produced by iMMC, MMC and MCX are shown in (d). The contour plot is limited to regions of the domain within 3 mm depth from the source due to higher stochastic noise in lower regions.

In Table 2, we summarize the numbers of nodes and elements for a side-by-side comparison between iMMC and DMMC. The mesh generation run-times (second) and the simulation speeds (photons/ms) are also reported in Table 2. Specifically, the mesh generation run-time for iMMC includes both the time for mesh generation using TetGen given the center-line information and that for labeling process using MATLAB.

**Table 2.** The performance comparison between implicit mesh-based Monte Carlo (iMMC) and dual-grid mesh-based Monte Carlo (DMMC) for a highly complex microvascular network. The run-time (second) for mesh generation and the simulation speed (photons/ms) are reported. For iMMC, the mesh generation run-time includes both tessellation using TetGen and edge-labeling process in MATLAB.

| Method | Node # | Element # | Meshing time (s) | Speed |
| --- | --- | --- | --- | --- |
| DMMC | 1,463,924 | 9,074,036 | 258.61 | 333 |
| iMMC | 6,850 | 44,209 | 26.20 | 966 |

## 4. Discussions and Conclusion

The meshes shown in Fig. 4 demonstrate not only a drastic reduction in element numbers from MMC to DMMC, as previously shown [17], but also that the hybrid shape representation used in iMMC achieves an additional reduction in mesh density and complexity in all 3 benchmarks. These results are further confirmed in Table 1: the number of elements in the iMMC mesh generated for benchmark B1 is 26,112× and 1,305× coarser than those of MMC and DMMC, respectively. For benchmarks B2 and B3, the element numbers for iMMC are 100-fold and 3-fold less, respectively, when compared to DMMC meshes, and an impressive over 2,500-fold reduction when compared to MMC. We want to mention that our current implement of e-iMMC has not yet supported feature-handling in adjacent elements, as shown in green in Fig 4(c). The development of a more rigorous simulation is underway.

The dramatically simplified mesh data results in improved simulation speed – from Table 1, a 4.8× to 5.4× speedup is achieved when comparing iMMC to MMC in all three benchmarks; when compared to DMMC, speed improvements of 7.1%, 7.1% and 9.3% are observed for B1, B2 and B3, respectively. The relatively small speedup over DMMC is due to the fact that both of them utilize coarse meshes; the computational overhead related to dense elements becomes relatively small compared to the necessary ray-tracing computation moving photons across tissue interfaces.

In Fig. 5, an excellent match can be observed between iMMC and DMMC simulation results for all benchmarks. The MMC results show minor mismatches when compared to those of iMMC and DMMC, especially near the light source. This has been previously reported in [17] as a result of insufficient output resolution in MMC. Fig. 5 provides strong evidence that iMMC is capable of achieving a similar level of accuracy as DMMC and MMC despite using only a fractional memory footprint for the mesh data. We want to point out that although iMMC eliminates the discretization errors associated with representing a curved surface feature with triangles, the accuracy of this approach relies on how closely the tissue structures resemble the assumed tubular, spherical or uniform layer feature shapes; when the actual anatomy deviates from these assumptions, the use of DMMC and MMC remains valid options; alternatively, introducing additional implicit parameters to support more complex shapes – for example, using a truncated cone instead of a cylinder to model vessels, or using spherical harmonics to represent non-spherical inclusions – could again make iMMC an effective and accurate solution.

The examples shown in Fig. 6 represent a highly sophisticated microvascular network in comparison to the benchmarks in Fig. 4 or other tissue models explored in earlier research [32]. In Figs. 6(c) and 6(d), we can visually observe that a considerable amount of voxels and tetrahedral elements are used to delineate the vessel boundaries in MCX and DMMC domain models. In comparison, the iMMC method in Fig. 6(b) significantly simplifies the domain complexity. As shown in Table 2, iMMC achieves over 213× and 205× reduction in node and element numbers, respectively, when compared to DMMC. Although DMMC does not introduce additional overhead by suppressing the refinement of tetrahedral elements within the same tissue region, it does require the creation of a triangular surface to discretize the complex shapes of dense networks of vessels with small radii; as a result, DMMC meshes contains nearly 1.5 million nodes – almost 99% of those nodes belong to vessel surfaces. The missing 1-voxel-with vessel branch in the DMMC mesh, indicated by the blue circle in Fig. 6(d), suggests that even higher surface mesh density may be necessary to accurately discretize such a complex domain using a tetrahedral/surface mesh representation. Unlike MMC/DMMC, the vessels in iMMC are represented by edges, which avoids dense tessellations at each vascular surface. The reduction in mesh density brings a significant speedup in mesh generation. In Table 2, a nearly 10× reduction in mesh generation run-time is achieved when comparing iMMC to DMMC. For iMMC, the actual run-time for mesh generation using TetGen is only 0.033s. Further acceleration of the labeling process is one of the further steps for iMMC code development.

Furthermore, due to the dramatic mesh simplification, iMMC improves the overall simulation speed (photons/ms) by 2.9-fold when compared to DMMC. The disproportionate relation between run-time reduction and reduction in mesh density can be attributed to the fact that standard scattering/absorption and boundary reflection – computations that are common for both iMMC and DMMC – become dominant as the mesh density reduces. Nevertheless, a 2.9-fold speedup for such a large-scale simulation is still quite impressive. We want to mention that we are currently working on implementing iMMC on the GPU [6] where high-speed memory is extremely limited and the host-device data transfer is expensive. In such cases, we anticipate higher speed gains when using iMMC on the GPU as a result of much fewer memory reading operations and better data caching.

In Fig. 7(d), an excellent match can be found between all three methods. Only in a small set of ultra-thin vessels, subtle differences can be observed due to the slight differences between rasterization approaches of these methods. The vessel structure in the iMMC result is nearly completely determined by vessel center-lines and radii input data, and does not have a dependency to rasterization resolution, which is the case when using tetrahedral and voxelated representations. However, as we highlighted in the red-circle region in Fig. 7(a), the lack of the ability to consider vessel shapes in the adjacent elements to a vessel-bearing element in our current implementation results in small artifacts in some large-diameter vessels. With further improvement of our software, we anticipate that such small errors will be removed in the future.

In summary, we have developed a significantly enhanced MMC algorithm that is tailored towards modeling extremely complex tissue structures – a highly challenging but increasingly important direction for biophotonic imaging. By using a hybrid approach that combines a coarse skeletal mesh and implicit shape parameters, iMMC reduces total memory usage by several hundred-to thousand-fold; this greatly simplified shape representation also results in about 5-fold speed improvement over conventional MMC in benchmark testing. In our three benchmarks, we also showed that edge-, node- and face-based iMMC simulations produced nearly identical solutions compared to standard MMC and DMMC. In addition, we have also successfully applied iMMC to model light transport in highly sophisticated microvascular networks. The node and element numbers are reduced by over 200-fold, which greatly reduces the memory usage when compared to the state-of-the-art DMMC method. The speed is also improved by 2.9-fold over DMMC due to the reduced overhead associated with simpler meshes. In the future, we will focus on implementing iMMC in our GPU-accelerated code [6] and supporting more advanced shape parameters, such as using truncated cones for vessel modeling. In addition, we will also explore further applications of iMMC in studying complex tissues, such as those of the lung and brain. The iMMC algorithm has been incorporated into our open-source MMC software and is freely available to the community at http://mcx.space/#mmc.

## Supporting information

Animation for Fig. 7a

## References

1. L. Wang, S. L. Jacques, and L. Zheng, “MCML – Monte Carlo modeling of light transport in multi-layered tissues,” Comput. methods programs biomedicine 47, 131–146 (1995).

2. C. Zhu and Q. Liu, “Review of Monte Carlo modeling of light transport in tissues,” J. Biomed. Opt. 18, 050902 (2013).

3. Q. Fang and D. A. Boas, “Monte Carlo simulation of photon migration in 3D turbid media accelerated by graphics processing units,” Opt. Express 17, 20178–20190 (2009).

4. N. Ren, J. Liang, X. Qu, J. Li, B. Lu, and J. Tian, “GPU-based Monte Carlo simulation for light propagation in complex heterogeneous tissues,” Opt. express 18, 6811–6823 (2010).

5. T. Young-Schultz, S. Brown, L. Lilge, and V. Betz, “FullMonteCUDA: a fast, flexible, and accurate GPU-accelerated Monte Carlo simulator for light propagation in turbid media,” Biomed. Opt. Express 10, 4711–4726 (2019).

6. Q. Fang and S. Yan, “Graphics processing unit-accelerated mesh-based Monte Carlo photon transport simulations,” J. biomedical optics 24, 115002 (2019).

7. M. A. Yücel, J. J. Selb, T. J. Huppert, M. A. Franceschini, and D. A. Boas, “Functional near infrared spectroscopy: Enabling routine functional brain imaging,” Curr. opinion biomedical engineering 4, 78–86 (2017).

8. Y. Yuan, P. Cassano, M. Pias, and Q. Fang, “Transcranial photobiomodulation with near-infrared light from childhood to elderliness: simulation of dosimetry,” Neurophotonics 7, 1–15 (2020).

9. P. Cassano, A. P. Tran, H. Katnani, B. S. Bleier, M. R. Hamblin, Y. Yuan, and Q. Fang, “Selective photobiomodulation for emotion regulation: model-based dosimetry study,” Neurophotonics 6, 1–12 (2019).

10. M. B. Robinson, S. A. Carp, A. Peruch, D. A. Boas, M. A. Franceschini, and S. Sakadžić, “Characterization of continuous wave ultrasound for acousto-optic modulated diffuse correlation spectroscopy (aom-dcs),” Biomed. Opt. Express 11, 3071–3090 (2020).

11. D. Hu, T. Sun, L. Yao, Z. Yang, A. Wang, and Y. Ying, “Monte carlo: A flexible and accurate technique for modeling light transport in food and agricultural products,” Trends Food Sci. Technol. (2020).

12. D. Wangpraseurt, S. You, F. Azam, G. Jacucci, O. Gaidarenko, M. Hildebrand, M. Kühl, A. G. Smith, M.P. Davey, A. Smith, D. D. Deheyn, S. Chen, and S. Vignolini, “Bionic 3d printed corals,” Nat. Commun. p. 1748 (2020).

13. D. A. Boas, J.P. Culver, J. J. Stott, and A. K. Dunn, “Three dimensional Monte Carlo code for photon migration through complex heterogeneous media including the adult human head,” Opt. express 10, 159–170 (2002).

14. S. Jacques and T. Li, “Monte Carlo simulations of light transport in 3D heterogenous tissues (mcxyz.c),” See http://omlc.org/software/mc/mcxyz/index.html [accessed 30.01. 2017] (2013).

15. S. Yan and Q. Fang, “Hybrid mesh and voxel based monte carlo algorithm for accurate and efficient photon transport modeling in complex bio-tissues,” Biomed. Opt. Express 11, 6262–6270 (2020).

16. Q. Fang, “Mesh-based Monte Carlo method using fast ray-tracing in Plücker coordinates,” Biomed. Opt. Express 1, 165–175 (2010).

17. S. Yan, A. P. Tran, and Q. Fang, “Dual-grid mesh-based Monte Carlo algorithm for efficient photon transport simulations in complex three-dimensional media,” J. Biomed. Opt. 24, 020503 (2019).

18. A. P. Tran, S. Yan, and Q. Fang, “Improving model-based functional near-infrared spectroscopy analysis using mesh-based anatomical and light-transport models,” Neurophotonics 7, 1–18 (2020).

19. Q. Fang and D. R. Kaeli, “Accelerating mesh-based Monte Carlo method on modern CPU architectures,” Biomed. Opt. Express 3, 3223–3230 (2012).

20. J. Cassidy, A. Nouri, V. Betz, and L. Lilge, “High-performance, robustly verified Monte Carlo simulation with FullMonte,” J. Biomed. Opt. 23, 085001 (2018).

21. C. J. Zoller, A. Hohmann, F. Forschum, S. Geiger, M. Geiger, T. P. Ertl, and A. Kienle, “Parallelized Monte Carlo software to efficiently simulate the light propagation in arbitrarily shaped objects and aligned scattering media,” J. Biomed. Opt. 23, 1–12–12 (2018).

22. J. Chen, Q. Fang, and X. Intes, “Mesh-based Monte Carlo method in time-domain widefield fluorescence molecular tomography,” J. Biomed. Opt. 17, 1–8 (2012).

23. R. Yao, X. Intes, and Q. Fang, “Generalized mesh-based Monte Carlo for wide-field illumination and detection via mesh retessellation,” Biomed. Opt. Express 7, 171–184 (2016).

24. A. A. Leino, A. Pulkkinen, and T. Tarvainen, “ValoMC: a Monte Carlo software and MATLAB toolbox for simulating light transport in biological tissue,” OSA Continuum 2, 957–972 (2019).

25. S. Powell and T. S. Leung, “Highly parallel monte-carlo simulations of the acousto-optic effect in heterogeneous turbid media,” J. Biomed. Opt. 17, 045002 (2012).

26. R. Yao, X. Intes, and Q. Fang, “Direct approach to compute Jacobians for diffuse optical tomography using perturbation Monte Carlo-based photon ‘replay’,” Biomed. Opt. Express 9, 4588–4603 (2018).

27. S. Yan, R. Yao, X. Intes, and Q. Fang, “Accelerating Monte Carlo modeling of structured-light-based diffuse optical imaging via “photon sharing”,” Opt. Lett. 45, 2842–2845 (2020).

28. M. Azimipour, M. Sheikhzadeh, R. Baumgartner, P. K. Cullen, F. J. Helmstetter, W.-J. Chang, and R. Pashaie, “Fluorescence laminar optical tomography for brain imaging: system implementation and performance evaluation,” J. Biomed. Opt. 22, 016003 (2017).

29. A. E. Hartinger, A. S. Nam, I. Chico-Calero, and B. J. Vakoc, “Monte Carlo modeling of angiographic optical coherence tomography,” Biomed. Opt. Express 5, 4338–4349 (2014).

30. C. Wang, L. Qiao, Z. Mao, Y. Cheng, and Z. Xu, “Reduced deep-tissue image degradation in three-dimensional multiphoton microscopy with concentric two-color two-photon fluorescence excitation,” JOSA B 25, 976–982 (2008).

31. X. Deng and M. Gu, “Penetration depth of single-, two-, and three-photon fluorescence microscopic imaging through human cortex structures: Monte Carlo simulation,” Appl. Opt. 42, 3321–3329 (2003).

32. L. Giannoni, F. Lange, and I. Tachtsidis, “Investigation of the quantification of hemoglobin and cytochrome-c-oxidase in the exposed cortex with near-infrared hyperspectral imaging: a simulation study,” J. Biomed. Opt. 25, 046001 (2020).

33. N. Ren, J. Liang, X. Qu, J. Li, B. Lu, and J. Tian, “GPU-based Monte Carlo simulation for light propagation in complex heterogeneous tissues,” Opt. Express 18, 6811–6823 (2010).

34. V. Periyasamy and M. Pramanik, “Monte Carlo simulation of light transport in turbid medium with embedded object – spherical, cylindrical, ellipsoidal, or cuboidal objects embedded within multilayered tissues,” J. biomedical optics 19, 045003 (2014).

35. Y. Zhang, B. Chen, D. Li, and G.-X. Wang, “Efficient and accurate simulation of light propagation in bio-tissues using the three-dimensional geometric Monte Carlo method,” Numer. Heat Transfer, Part A: Appl. 68, 827–846 (2015).

36. V. Periyasamy and M. Pramanik, “Importance sampling-based Monte Carlo simulation of time-domain optical coherence tomography with embedded objects,” Appl. optics 55, 2921–2929 (2016).

37. L. Gagnon, S. Sakadžić, F. Lesage, J. J. Musacchia, J. Lefebvre, Q. Fang, M. A. Yücel, K. C. Evans, E. T. Mandeville, J. Cohen-Adad, J. R. Polimeni, M. A. Yaseen, E. H. Lo, D. N. Greve, R. B. Buxton, A. M. Dale, A. Devor, and D. A. Boas, “Quantifying the microvascular origin of bold-fmri from first principles with two-photon microscopy and an oxygen-sensitive nanoprobe,” J. Neurosci. 35, 3663–3675 (2015).

38. H. Si, “TetGen, a Delaunay-based quality tetrahedral mesh generator,” ACM Transactions on Math. Softw. (TOMS) 41, 1–36 (2015).

39. G. Tetteh, V. Efremov, N. D. Forkert, M. Schneider, J. Kirschke, B. Weber, C. Zimmer, M. Piraud, and B. H. Menze, “Deepvesselnet: Vessel segmentation, centerline prediction, and bifurcation detection in 3-D angiographic volumes,” arXiv preprint arXiv:1803.09340 (2018).

40. Q. Fang and D. A. Boas, “Tetrahedral mesh generation from volumetric binary and grayscale images,” in 2009 IEEE International Symposium on Biomedical Imaging: From Nano to Macro, (IEEE, 2009), pp. 1142–1145.

