## Supplementary figures and images for "Light transport modeling in highly complex tissues using implicit mesh-based Monte Carlo algorithm"

### Animation for Fig. 7a

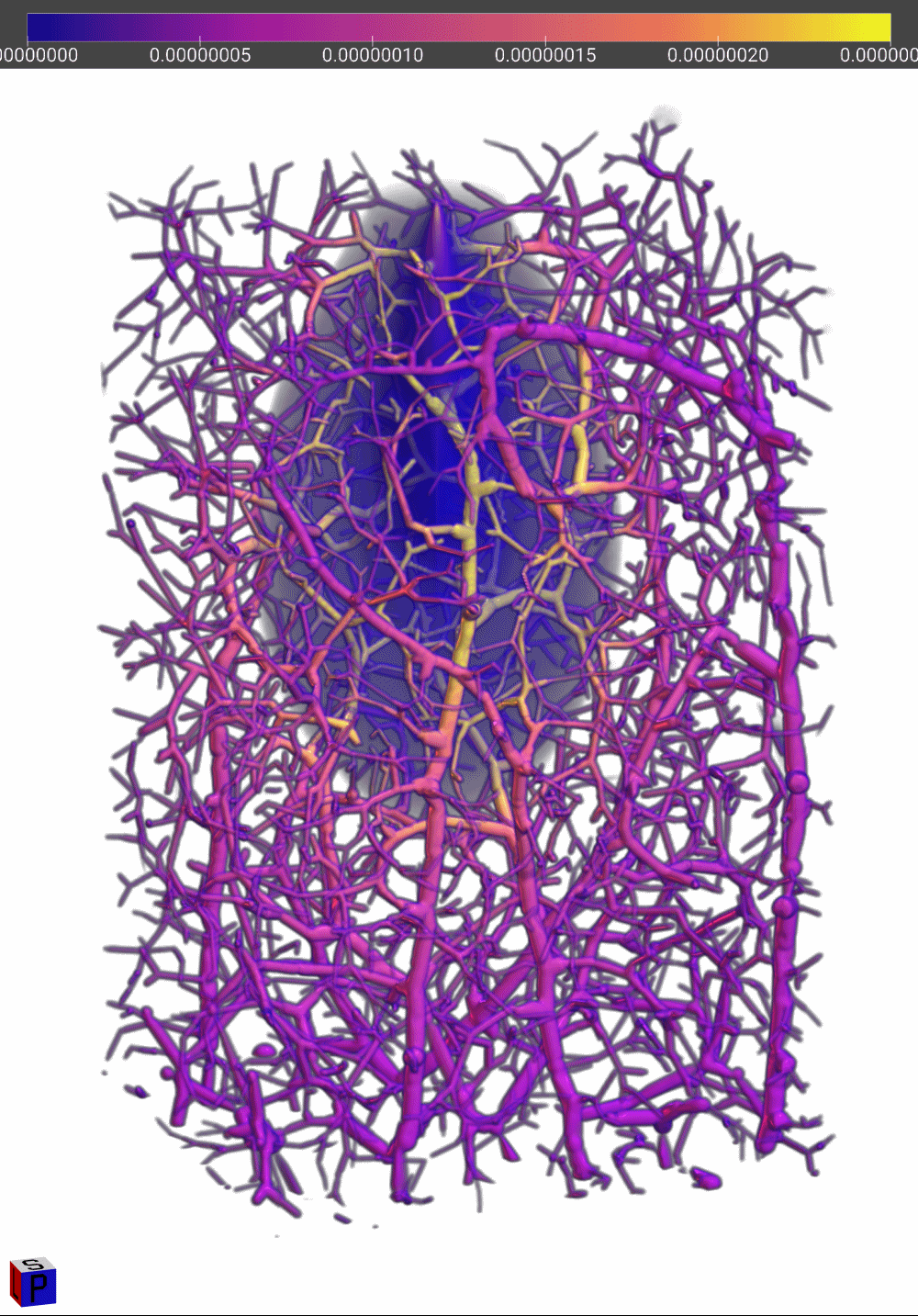
